# Repurpose of Ritlecitinib Reduces Neointima Formation and Enhances Endothelial Recovery Following Vascular Injury

**DOI:** 10.1101/2025.09.23.678172

**Authors:** Dunpeng Cai, Yung-Chun Wang, Lindsey Saint, John Markley, Shi-You Chen

## Abstract

**Background:** Neointimal hyperplasia after vascular injury reflects excess smooth muscle cell (SMC) proliferation with impaired endothelial recovery. We tested the hypothesis that selective Janus kinase 3 (JAK3) inhibition with the FDA-approved drug Ritlecitinib limits neointima while accelerating reendothelialization.

**Methods:** We used mouse carotid wire injury and a human internal mammary artery (IMA) xenograft model to evaluate effects of genetic JAK3 loss in SMCs or oral Ritlecitinib in vascular remodeling. Mechanistic studies in SMCs assessed JNK/c-Jun signaling and its regulation of thrombospondin-1 (TSP-1) and vascular endothelial growth factor A (VEGF-A) using RNA-seq and CUT&RUN. Rescue experiments tested whether VEGF-A neutralization or exogenous TSP-1 reversed the antineointimal effects. Patient-derived IMA SMCs were analyzed for variabilities in drug responses.

**Results:** Genetic JAK3 loss in SMCs or oral Ritlecitinib reduced neointimal area and intima/media ratio and increased luminal CD31+ coverage in both models. In SMCs, Ritlecitinib suppressed JNK/c-Jun signaling, downregulated TSP-1, and restored VEGF-A, shifting the milieu toward endothelial regeneration. RNA-seq and CUT&RUN corroborated JNK/c-Jun– dependent control of TSP-1 and VEGFA. VEGF-A neutralization or exogenous TSP-1 abrogated the anti-neointimal phenotype. In patient-derived IMA SMCs, variable drug response was linked to the JAK3 Pro893Asn variant and a VEGFA promoter SNP.

**Conclusion:** Selective JAK3 inhibition by Ritlecitinib simultaneously restrains SMC proliferation and promotes endothelial repair through modulating TSP-1 and VEGF-A activities. Ritlecitinib merits clinical evaluation as a precision therapy to prevent restenosis, with genetic stratification to identify responders.

## INTRODUCTION

The integrity of the vascular endothelium is essential for maintaining the overall health of blood vessels^1-6^. Endothelial cells (ECs) regulate critical vascular processes, such as maintaining vascular tone, controlling inflammation, preventing thrombosis, and facilitating angiogenesis^7-13^. When vascular interventions like balloon angioplasty or stent implantation are performed to treat atherosclerotic obstructions, they often damage the endothelium^14-21^. This damage disrupts normal vascular functions and leaves the vessel wall vulnerable to complications such as thrombosis^18,22^. To restore a functional endothelial barrier, EC proliferation is vital, and factors such as shear stress and the inherent properties of ECs determine how effectively this recovery occurs^23-26^. Vascular smooth muscle cells (SMCs), located beneath the endothelium, also play an important role in promoting EC recovery^27-30^. SMCs release growth factors, including fibroblast growth factor (FGF)^31^ and vascular endothelial growth factor (VEGF)^32^, which support EC proliferation and help reestablish an intact endothelium. However, mechanisms through which SMCs influence EC repair following vascular injury remain poorly understood.

Neointimal hyperplasia, characterized by excessive proliferation of SMCs and impaired EC recovery, is a key contributor to restenosis following cardiovascular interventions such as percutaneous coronary intervention (PCI) or bypass grafting^33,34^. Despite advancements in the development of drug-eluting stents and anti-proliferative therapies, restenosis remains a significant challenge, emphasizing the need for new therapeutic approaches^35^. Although drug-releasing stents, such as those coated with paclitaxel or sirolimus, have been effective in preventing restenosis by delivering localized therapy, they have also shown delayed vascular healing and “late catch-up” effects, where neointimal proliferation resumes after the initial inhibition^36^. This limitation points to the necessity of systemic adjunctive therapies to maintain neointimal inhibition and reduce the risk of restenosis over time^37^. Current systemic therapies, such as lipid-lowering agents, antioxidants, and antithrombotic drugs, have shown limited success in preventing restenosis^38^. Furthermore, pharmacokinetic studies of oral agents like sirolimus have revealed substantial interindividual variability, which complicates their therapeutic use^39^. Thus, there is a critical need to develop oral pharmacologic agents that can effectively prevent neointimal hyperplasia while promoting endothelial recovery in a controlled manner.

Neointimal hyperplasia is primarily driven by increased SMC proliferation alongside EC proliferation, disrupting the balance necessary for effective vessel healing^40^. Thus, therapies that can simultaneously inhibit SMC proliferation while promoting EC recovery are urgently needed to improve vascular repair outcomes. JAK3, a nonreceptor tyrosine kinase, mediates cytokine signaling through receptors containing the common γ chain and influences various cell types in both physiological and pathological contexts^41^. Notably, the absence of JAK3 impairs the development of immune cells, such as natural killer (NK) cells, T cells, and B cells, and affects the proliferation of lung epithelial cells^42^. Our previous research demonstrated that JAK3 is essential for SMC proliferation^43^. In the current study, we found that JAK3 deficiency in SMCs enhances EC proliferation through increased secretion of VEGF-A while reducing secretion of thrombospondin-1 (TSP-1). By building on prior findings, this study further investigates the potential of JAK3 as a therapeutic target for managing neointimal hyperplasia and promoting vascular repair in human arteries.

Ritlecitinib, a selective JAK3 inhibitor, was originally developed for the treatment of alopecia areata, an autoimmune disease characterized by hair loss^44^. Given the established role of JAK3 in modulating immune responses, as well as its involvement in SMC proliferation, we hypothesized that inhibiting JAK3 with Ritlecitinib could effectively reduce neointima formation while promoting endothelial healing. Thus, we evaluated the efficacy of Ritlecitinib in preclinical models of arterial injury of human arteries, with the ultimate goal of repurposing this FDA-approved drug as a novel therapeutic strategy for vascular disease, particularly for preventing restenosis in high-risk patients through personalized medicine. Interestingly, through large-scale preclinical testing with Ritlecitinib in human internal mammary artery (IMA) balloon injury models, we found that the effect of balloon angioplasty injury varied significantly among different patients’ IMAs. Specifically, while most patients responded well to Ritlecitinib treatment with substantial neointima reduction, a subset of patients exhibited minimal response. To explore the underlying mechanisms of this variability, SMCs from both responders and non-responders were isolated and analyzed. Genetic analyses revealed that a mutation in the JAK3 protein (Pro893Asn) and a single nucleotide polymorphism (SNP) in the VEGF-A promoter region were associated with the reduced efficacy of Ritlecitinib in non-responders. These findings underscore the importance of a precision medicine approach, leading to personalized medical interventions through tailoring therapeutic strategies to individual patient’s genetic characteristics.

## METHODS

### Data Availability

The data underlying this article are available in the article and in its Supplemental Material.

### Animal Models and Carotid Artery Wire Injury

Male ApoE-/-mice on C57BL/6 background (8–10 weeks old; Jackson Labs) were used for the wire injury induced neointima model. To specifically ablate JAK3 in VSMCs, we bred Myh11-Cre-ERT2; eYFPfl/fl; JAK3fl/fl mice into the ApoE-/-background. Tamoxifen (75 mg/kg i.p. for 5 consecutive days) was given at 6 weeks of age to induce Cre recombination and knock out JAK3 in smooth muscle cells (Myh11Cre; eYFPfl/fl; JAK3fl/fl;ApoE-/-). Littermate Myh11Cre; eYFPfl/fl; ApoE-/-mice served as controls. All mice were fed a chow diet. At 10 weeks of age, mice underwent left common carotid artery endoluminal denudation injury under isoflurane anesthesia as we previously reported^43^. Briefly, the left external carotid artery was ligated and a small arteriotomy made. A 0.38 mm diameter flexible guidewire (Cook Medical, G02426) was inserted via the external carotid into the common carotid artery and passed toward the aortic arch, then withdrawn with rotation. This wire passage was repeated 6 times to thoroughly denude endothelium. The external carotid was then permanently ligated and blood flow guided to the internal common carotid. Buprenorphine was given for analgesia. Sham surgery (vessel exposure without wire injury) was performed on the right carotid artery. Mice were euthanized at 4 weeks post-injury for tissue analysis, a timeframe sufficient for neointima formation. For Ritlecitinib (T5382-100MG; Sigma) administration, the drug was formulated fresh in 0.5% methylcellulose and administered by oral gavage at 50 mg/kg/day, starting on the day of carotid injury and continued daily until sacrifice. This dose was based on pilot tolerability and achieves effective JAK3 inhibition in vivo. Control mice received vehicle. Mice were monitored daily for health and body weight.

### Human Artery Injury and Transplant Model

Surplus segments of internal mammary artery (IMA) were obtained from coronary artery bypass graft (CABG) patients (age 50–75, both sexes) at the time of surgical operation under an approved IRB protocol (IRB # 2026026) with informed consent. Within 1 hour of harvest, peri-adventitial fat was trimmed, and each IMA segment (∼1.5–2 cm length) was subjected to endothelial injury using a 2-Fr Fogarty balloon catheter^45^. The catheter was inserted and inflated to gently appose the vessel wall, then passed through the lumen three times to denude endothelium at the pressure of 1.5 atm. To modulate JAK3 expression in human vessels, IMA segments were incubated with adenoviruses encoding either a JAK3-targeted shRNA (Ad-shJAK3) or a GFP control (Ad-GFP)^43^. High-titer adenoviral stocks (∼10^9 PFU/mL) were delivered into the lumen of each IMA segment ex vivo, i.e., the segment was cannulated with an 24G catheter, and the adenovirus solution was infused to distend the lumen with the ends clamped. After 30 minutes of incubation at 37°C, allowing transduction to the arterial media, the virus was flushed out and segments were cultured overnight in DMEM with 2% FBS to permit gene knockdown. This procedure achieved efficient transgene expression in the vessel wall as confirmed by GFP visualization. The following day the human IMA segments with injury and adenoviral transduction were implanted into immunodeficient NOD/SCID-IL2Rγ null (NSG) mice (The Jackson Laboratory) to allow in vivo remodeling as we reported previously ^46^. Briefly, 10–12 weeks old mice (both sexes) were anesthetized with 2–3% isoflurane in oxygen, and a midline laparotomy was performed to expose the abdominal aorta. A 5–7 mm segment of the abdominal aorta was excised and replaced with the denuded human IMA segment using end-to-end anastomosis over polyethylene cuffs secured with 11-0 nylon sutures under a surgical microscope. This xenograft model permits human arterial tissue to undergo neointimal hyperplasia under in vivo conditions of host cell infiltration and low shear stress, in the absence of direct human blood perfusion.

NSG mice carrying IMA grafts were treated daily by oral gavage with Ritlecitinib (50 mg/kg, as in the murine injury model) or vehicle to evaluate the effect of drug on human vessel remodeling. In separate experimental groups, additional interventions were applied: bevacizumab (5 mg/kg, i.v, twice weekly; Avastin, Genentech) was used to block VEGF-A signaling, while recombinant human thrombospondin-1 (TSP-1, 5 μg/graft; R&D Systems, #3074-TH) was delivered by i.v injection on days 0 and 7 to enhance TSP-1 activity. Grafts were harvested 14 days post-implantation for histological and molecular analyses, a time point established in pilot studies as sufficient for the development of neointima and early endothelial recovery. NSG mice were maintained on drinking water containing enrofloxacin for infection prophylaxis and monitored daily for post-surgical recovery and signs of graft-related complications.

## Statistical Analysis

Data are presented as mean ± SEM. Statistical analyses were performed using GraphPad Prism 9. For comparisons between two groups, an unpaired two-tailed Student’s t-test was used (with Welch’s correction if variance was unequal). For comparisons among multiple groups (≥3), one-way or two-way ANOVA was applied as appropriate, followed by Tukey’s post hoc test for pairwise contrasts. The specific tests used for each experiment are indicated in the figure legends. For non-parametric data (e.g. histological scoring), a Mann–Whitney U test or Kruskal– Wallis test was used. P < 0.05 was considered statistically significant. In figures, significance is annotated as P<0.05, *P<0.01, and **P<0.001. No samples or animals were excluded from analysis unless pre-established criteria for technical failures were met (e.g., graft thrombosis).

All data met assumptions of the statistical tests used (normality and homoscedasticity when required, tested via Shapiro–Wilk and Levene’s tests). The investigators had no control over randomization for human samples (which were de-identified), but treatment conditions were randomized in cell culture experiments. All statistical tests were two-sided.

## RESULTS

### Ritlecitinib Selectively Blocks JAK3 Activity and Suppresses SMC Proliferation

Since our previous studies have shown that JAK3 promotes SMC proliferation, we sought to evaluate the specificity and efficacy of Ritlecitinib, an FDA-approved drug, in inhibiting JAK3 signaling in SMCs. SMCs isolated from human IMAs were treated with various JAK inhibitors, including Ruxolitinib, Baricitinib, Fedratinib, and Ritlecitinib. Western blot analysis of JAK3 phosphorylation revealed that Ritlecitinib completely suppressed JAK3 activation, whereas the other inhibitors, primarily targeting JAK1 or JAK2, showed minimal or partial effects (Fig. 1A-B). These data indicate that Ritlecitinib is highly selective for JAK3 in SMCs, effectively blocking JAK3 kinase activity. In line with this selectivity, Ritlecitinib also disrupted JAK3 downstream signaling pathways. Platelet-derived growth factor-BB (PDGF-BB) stimulation of SMCs led to the activation of JAK3 and its downstream JNK and STAT3 pathways. Ritlecitinib markedly reduced the phosphorylation of key signaling proteins in these pathways (Fig. 1C), demonstrating that it effectively blocks JAK3-dependent downstream signals. Functionally, EdU incorporation assay showed that PDGF-BB induced robust SMC proliferation in control conditions, whereas Ritlecitinib-treated cells displayed far fewer EdU-positive nuclei (Fig. 1D). These results establish that Ritlecitinib selectively targets JAK3 in vascular cells and effectively curtails pathological SMC proliferation.

**Fig 1.**
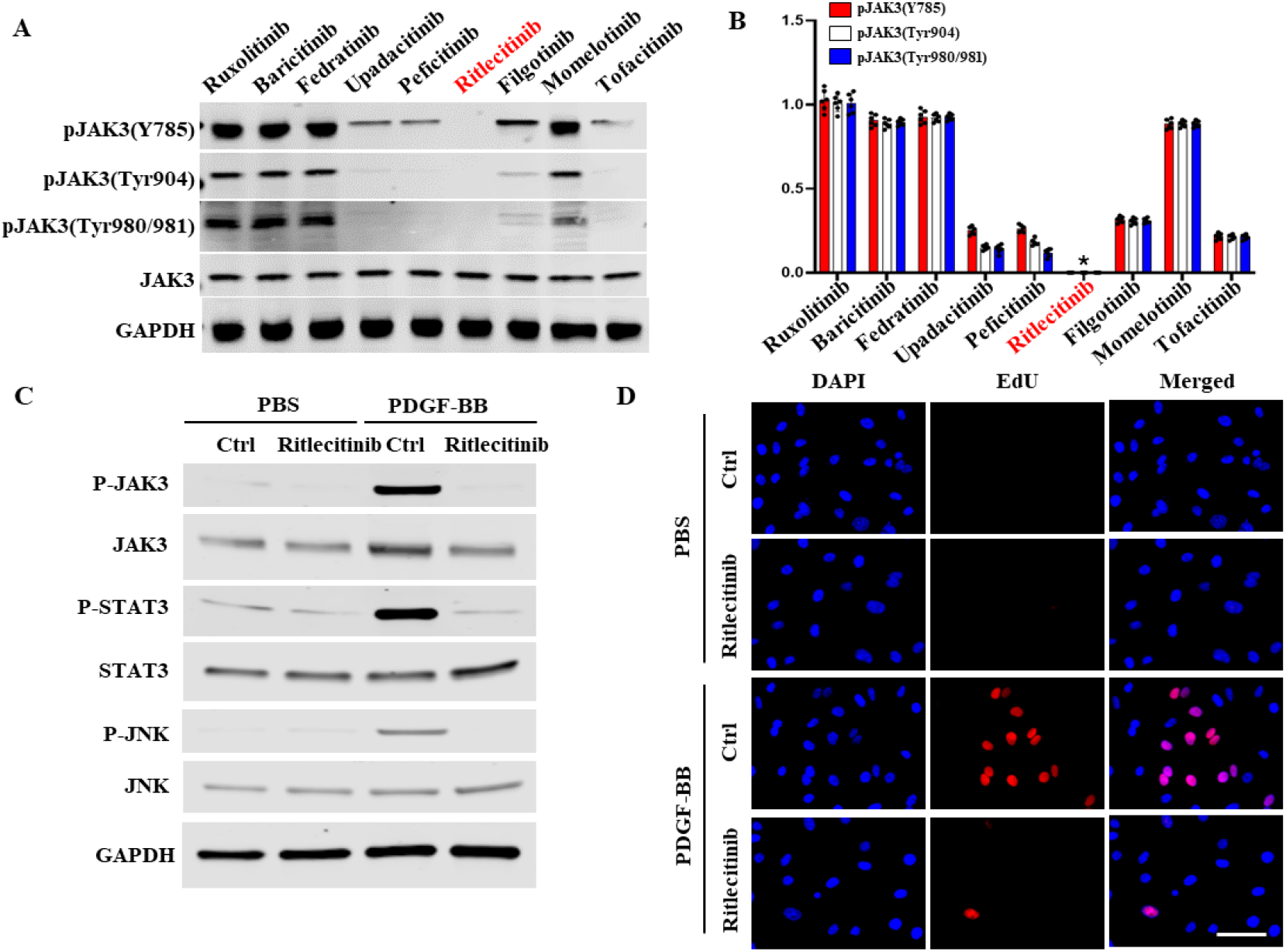
Ritlecitinib selectively blocks JAK3 activities and suppresses SMC proliferation. Human SMCs cultured from IMA segments leftover from CABG surgery were treated with various JAK inhibitors for 6 hours. Proteins were then extracted for analyses. **A**, Among many JAK3 inhibitors, only Ritlecitinib selectively inhibited JAK3 phosphorylation. Ritlecitinib completely blocked phosphorylation of JAK3 at multiple sites, while other JAK inhibitors exhibited minimal or partial effects. **B**, Quantification of JAK3 phosphorylation levels from panel A, confirming the significant reduction in JAK3 phosphorylation by Ritlecitinib. *P<0.05 vs. other inhibitors. **C**, Ritlecitinib effectively blocks JAK3 downstream signaling pathways in PDGF-BB-treated human SMCs as shown by the reduced phosphorylation of key signaling proteins. **D**, Ritlecitinib inhibited SMC proliferation induced by PDGF-BB as shown by EdU staining (red). DAPI stains nuclei. Ritlecitinib treatment significantly diminished the EdU+ SMCs, indicating reduced SMC proliferation.

### Ritlecitinib Suppresses Neointima Formation and Accelerates Reendothelialization in Injured Mouse Arteries

We then investigated the therapeutic effects of Ritlecitinib in a murine model of arterial injury, which recapitulates neointimal hyperplasia and endothelial regeneration seen in clinical restenosis. ApoE-/-mice underwent carotid artery wire denudation and were treated daily with either saline (control) or Ritlecitinib (50 mg/kg, oral gavage) starting at day one post-injury.

Histological assessment at 21 days post-injury demonstrated a dramatic reduction in neointimal hyperplasia in Ritlecitinib-treated arteries compared to controls (Fig. 2A). Morphometric quantification showed that the neointimal area and the intima-to-media area ratio (I/M ratio) were significantly lower in Ritlecitinib-treated arteries as compared to the saline-treated ones (Fig. 2B-C). Importantly, consistent with its anti-proliferative effects, Ritlecitinib treatment markedly reduced SMC proliferation in the injured arterial wall, as evidenced by diminished EdU labeling in SMCs (Supplemental Fig. 1A). These results indicate that Ritlecitinib attenuates the pathological accumulation of SMCs in the intima following injury.

**Fig 2.**
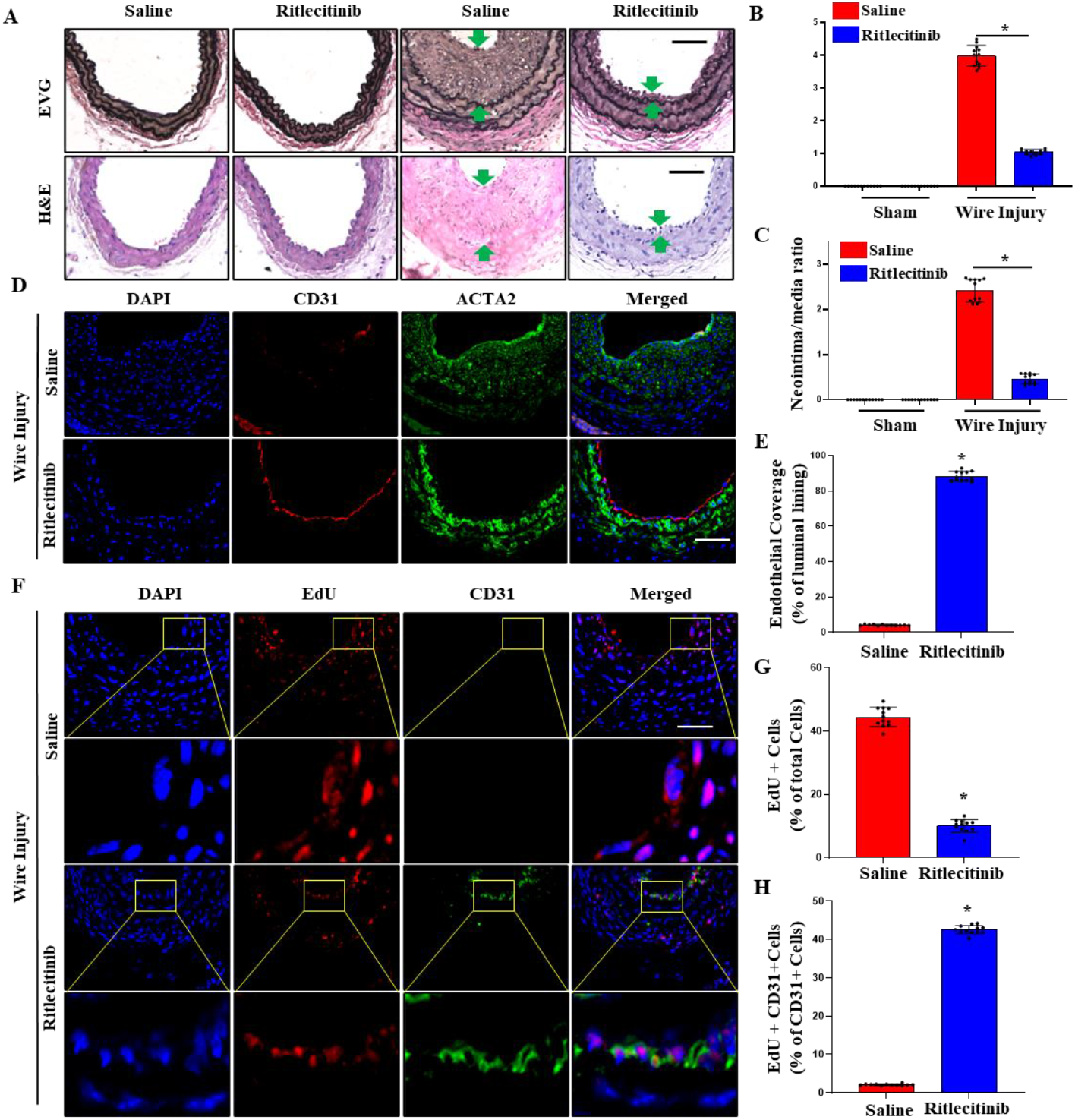
Ritlecitinib suppresses neointima formation and accelerates reendothelialization in wire-injured mouse arteries. ApoE-/-mouse carotid arteries were denuded by wire injury. The mice were administered with either saline or Ritlecitinib (50 mg/kg daily) starting from day 1 following the injury. The injured arteries were collected at 21 days post-injury for analyses, with contralateral uninjured arteries as controls (sham). **A**, Ritlecitinib blocked the injury-induced neointima formation, as shown by Elastica van Gieson (EVG) and H&E staining. Scale bar: 50 μm. **B-C**, Quantification of neointima areas (B) and intima/media ratios (C). *P<0.05 vs. saline-treated arteries, n=12. **D**, Ritlecitinib promoted reendothelialization as shown by CD31 immunostaining (red) in the lumen of injured arteries. The media layer was stained with ACTA2. Scale bar: 50 μm. **E**, Quantification of the endothelial coverage by measuring the percentage of lumen area lined by CD31+ cells. *P<0.05 vs. saline-treated arteries, n=12. **F**, Ritlecitinib promoted endothelial (CD31+) cell proliferation in injured arteries as shown by EdU staining (red) in the intima layer. Scale bar: 50 μm. **G**, Quantification of EdU+ cells relative to total CD31+ cells. *P<0.05 vs. saline-treated arteries, n=12. **H**, Quantification of EdU+CD31+ cells relative to total CD31+ cells. *P<0.05 vs. saline-treated arteries, n=12.

In addition to suppressing neointima formation, Ritlecitinib markedly enhanced endothelial recovery in the injured arteries. Immunostaining of CD31, an endothelial cell marker, demonstrated a substantially greater reendothelialization in Ritlecitinib-treated carotid arteries, as shown by CD31-positive cells forming a nearly continuous lining on the luminal surface in the Ritlecitinib group, whereas saline-treated arteries showed large denuded areas (Fig. 2D). The percentage of the lumen covered by CD31+ endothelium was also significantly higher with Ritlecitinib treatment (Fig. 2E). To determine if the improved endothelial coverage was due to increased endothelial cell proliferation, we performed EdU incorporation assays in vivo. Ritlecitinib-treated arteries exhibited a higher density of EdU-positive cells within the intima, many of which co-localized with CD31, indicating proliferating endothelial cells (Fig. 2F). Although the proportion of EdU+ cells among total intimal cells decreased in the Ritlecitinib group (Fig. 2G), due to inhibition of SMC proliferation, the percentage of EdU+ cells that were CD31+ endothelial cells were significantly increased as compared to controls (Fig. 2H). These data indicate that Ritlecitinib fosters a favorable EC-regenerating environment after vascular injury by inhibiting destructive neointimal SMC proliferation while promoting the regeneration of the endothelial layer. The dual action of Ritlecitinib observed in this model, i.e., limiting neointimal growth while expediting reendothelialization, highlights its promise as a therapeutic agent for preventing restenosis.

### JAK3 Knockdown Blocks Neointima Formation and Promotes Reendothelialization in Human Arteries

To translate these findings to human vascular tissue, we examined the impact of JAK3 suppression in an ex vivo human artery injury model. Segments of human IMA obtained from CABG surgery were subjected to deliberate endothelial injury using a balloon catheter and then transduced with adenoviral vector encoding JAK3 shRNA (Ad-shJAK3) or with control adenovirus (Ad-GFP). The arteries were then transplanted to abdominal aortas of immunodeficient NSG mice to allow healing and remodeling for 21 days. Histological analysis revealed that JAK3 knockdown profoundly attenuated neointimal hyperplasia in human arteries. Injured IMAs receiving Ad-shJAK3 showed a thin neointimal layer compared to the substantial neointima observed in control arteries (Fig. 3A). Morphometric quantification confirmed the significant decrease in neointimal area (Fig. 3B) and intima/media ratio (Fig. 3C) in Ad-shJAK3 group relative to controls, underscoring the role of JAK3 in driving human arterial intimal thickening. Concomitant with the reduced neointimal formation, JAK3 knockdown significantly improved endothelial recovery in the injured human arteries. CD31 immunostaining showed that Ad-shJAK3–treated IMAs had much more endothelial lining on the luminal surface than control arteries, which remained partially denuded (Fig. 3D). Quantitative analysis demonstrated that the percentage of lumen with CD31+ endothelial coverage was significantly higher in JAK3-deficient arteries (Fig. 3E). These results indicate that inhibiting JAK3 not only curbs pathological neointimal growth but also facilitates regenerative reendothelialization in human arteries.

**Fig 3.**
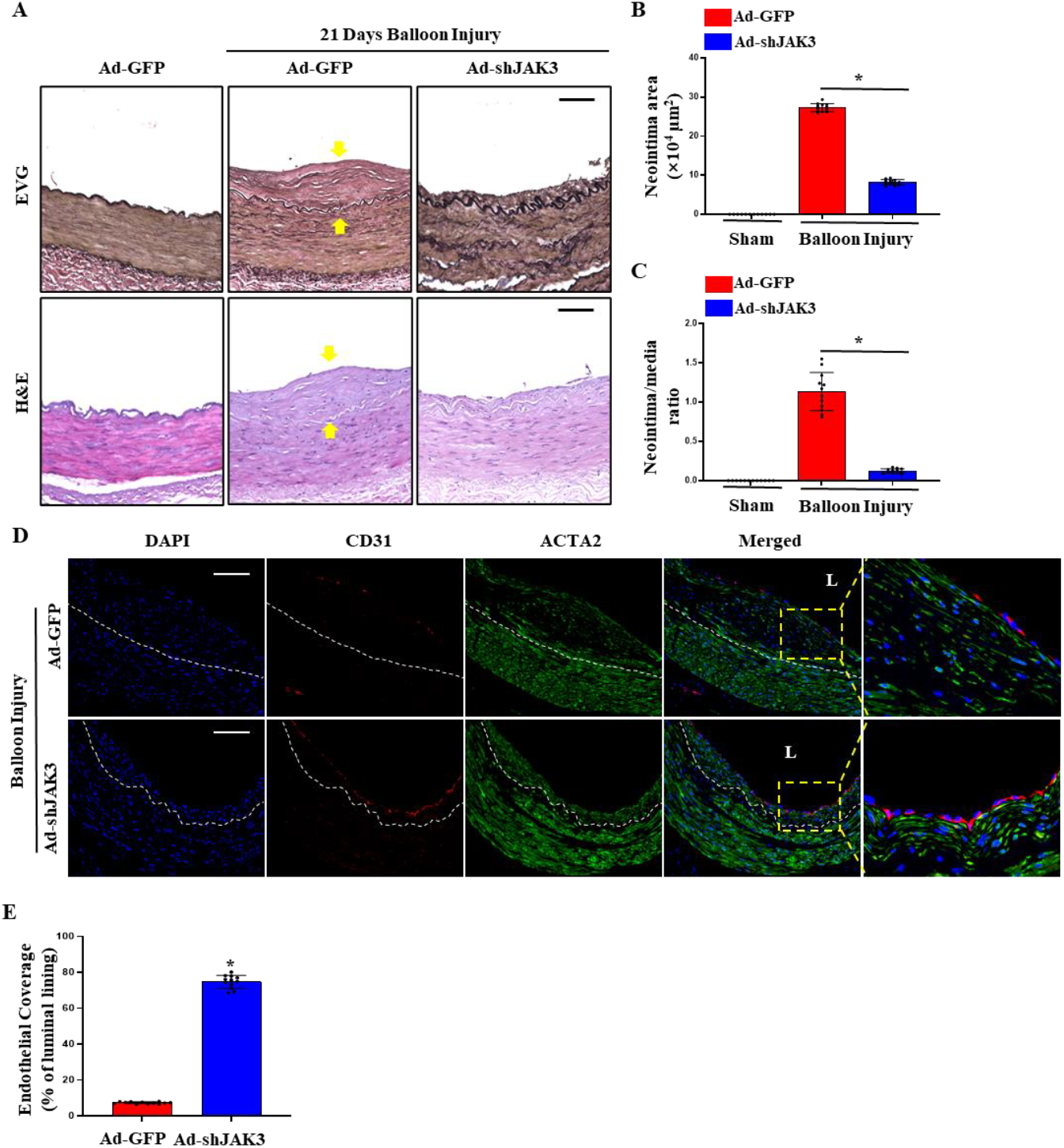
Knockdown of JAK3 blocked neointima formation while promoting re-endothelization in human arteries. Human IMA segments from CABG procedure were subjected to balloon injury to denude the endothelium. Ad-shJAK3 or Ad-GFP (control) was transduced in the lumen to knock down JAK3 in artery media. The IMAs were then transplanted to the abdominal aortas of immune-deficient NSG mice via end-to-end anastomosis. The IMAs were collected at 21 days post-injury. **A**, JAK3 knockdown diminished injury-induced neointima formation in human IMAs, as shown by Elastica van Gieson (EVG) and H&E staining. Scale bar: 100 μm. **B-C**, Quantification of neointima areas (B) and neointima/media ratios (C). *P<0.05 vs. Ad-GFP-transduced IMAs, n=12. **D**, JAK3 knockdown promoted reendothelialization, as shown by CD31 immunostaining (red) in the lumen of injured IMAs. The media and neointima areas were stained with ACTA2 (green). Scale bar: 100 μm. **E**, Quantification of the endothelial coverage by measuring the percentage of lumen area lined by CD31+ cells. *P<0.05 vs. Ad-GFP-transduced IMAs, n=12.

### Ritlecitinib Reduces Neointimal Hyperplasia and Promotes Endothelial Repair in Injured Human IMAs

Having demonstrated the benefits of JAK3 knockdown in human arteries, we next tested whether pharmacological JAK3 inhibition with Ritlecitinib could improve outcomes in injured human vessels. Balloon-injured human IMA segments were transplanted into NSG mice and treated systemically with either saline or Ritlecitinib (50 mg/kg daily) starting the day after injury. After 21 days, Ritlecitinib-treated human IMAs showed strikingly less neointimal hyperplasia than saline-treated controls. EVG and H&E staining revealed that the neointimal layer was thinner and contained fewer cells in the Ritlecitinib-treated group, whereas control arteries developed a robust neointima (Fig. 4A). Indeed, Ritlecitinib caused significant reduction in neointima area (Fig. 4B) and the intima/media ratio (Fig. 4C). The beneficial effects of Ritlecitinib in human arteries were associated with a reduction in SMC proliferation within the neointima. Co-staining of EdU with ACTA2 on IMA cross-sections revealed that Ritlecitinib-treated vessels had far fewer proliferating SMCs in the intima and media layers compared to controls (Supplemental Fig. 1B).

**Fig 4.**
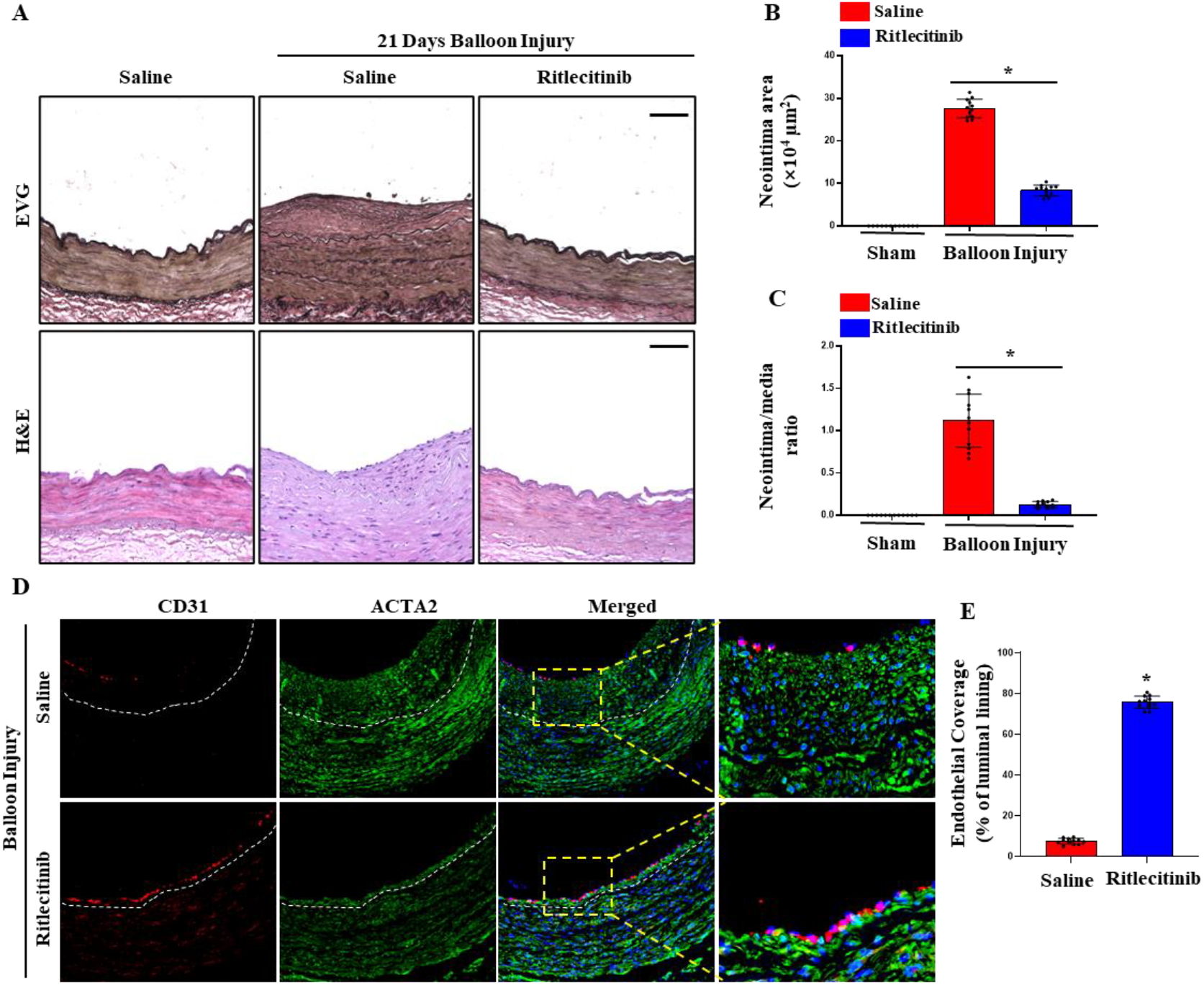
Ritlecitinib suppresses neointima formation and promotes reendothelialization in injured human arteries. Human IMA segments from CABG procedures were subjected to balloon injury to disrupt the endothelium. The injured IMAs were then transplanted to abdominal aortas of immune-deficient NSG mice. Saline or Ritlecitinib (50 mg/kg daily) was administered starting on day 1 post-transplantation. 21 days later, the IMAs were collected for analysis. **A**, Ritlecitinib significantly reduced neointima formation in injured human IMAs as demonstrated by Elastica van Gieson (EVG) and H&E staining. Scale bar: 100 μm. **B-C**, Quantification of neointima areas (B) and intima/media ratios (C). *P<0.05 vs. saline-treated IMAs, n=12. **D**, Ritlecitinib promoted reendothelialization in injured IMAs as shown by CD31 immunostaining (red). The media layer was stained with ACTA2. Scale bar: 100 μm. **E**, Quantification of the endothelial coverage by measuring the percentage of lumen area lined by CD31+ cells. *P<0.05 vs. saline-treated IMAs, n=12.

More importantly, Ritlecitinib markedly enhanced reendothelialization in the injured human arteries, as shown by the higher density of CD31+ ECs lining the lumen of Ritlecitinib-treated IMAs compared to controls (Fig. 4D). The percentage of the lumen covered by CD31+ endothelium was significantly greater in the Ritlecitinib group (Fig. 4E), reflecting improved endothelial healing. These findings mirror those observed in the mouse model, highlighting the translational potential of Ritlecitinib for preventing restenosis in human arteries.

Since targeting JAK3, either by genetic knockdown or pharmacological inhibition, leads to dual benefits: suppression of neointimal SMC proliferation and acceleration of endothelial repair, we sought to determine if these two effects are independent or linked events. Thus, we performed wire injury in carotid arteries of SMC-specific JAK3 deficient mice in ApoE-/-background (Myh11-Cre; JAK3fl/fl; ApoE-/-or JAK3smc-/-). The injury resulted in dramatically less neointima formation in JAK3smc-/-arteries compared to wild-type controls (Myh11-Cre;ApoE-/-mice), as shown by significantly smaller neointimal areas and lower intima/media ratios (Supplemental Fig. 2, A-C). Importantly, JAK3smc-/-mice also exhibited enhanced endothelial repair, with greater luminal CD31+ cell coverage and increased proliferation of endothelial cells relative to controls (Supplemental Fig. 2, D-H). These results indicated that neointimal SMC-produced JAK3 blocks EC proliferation and consequently endothelium recovery from the injury, and that the accelerated endothelial repair due to targeting JAK3 resulted from suppression of neointima.

### JAK3 Inhibition Modulates VEGF-A and TSP-1 Expression via the JNK/c-Jun Pathway

To elucidate the molecular mechanism by which JAK3 inhibition curbs neointimal hyperplasia which further promotes reendothelialization, we conducted transcriptomic and signaling analyses and identified two key counter-regulatory factors, i.e., vascular endothelial growth factor A (VEGF-A), a pro-endothelial regenerative factor, and thrombospondin-1 (TSP-1), an anti-angiogenic and pro-inflammatory matricellular protein. Bulk RNA sequencing (RNA-seq) of wire-injured mouse carotids (day 7 post-injury) revealed that Ritlecitinib treatment induced widespread changes in gene expression. A volcano plot of differentially expressed genes showed numerous transcripts significantly upregulated or downregulated by Ritlecitinib compared to saline treatment (Fig. 5A). Gene set enrichment analysis (GSEA) highlighted that genes in the JNK/c-Jun signaling pathway were among the most significantly affected. The enrichment plot demonstrated a strong negative enrichment score for JNK/c-Jun pathway gene sets in

**Fig 5.**
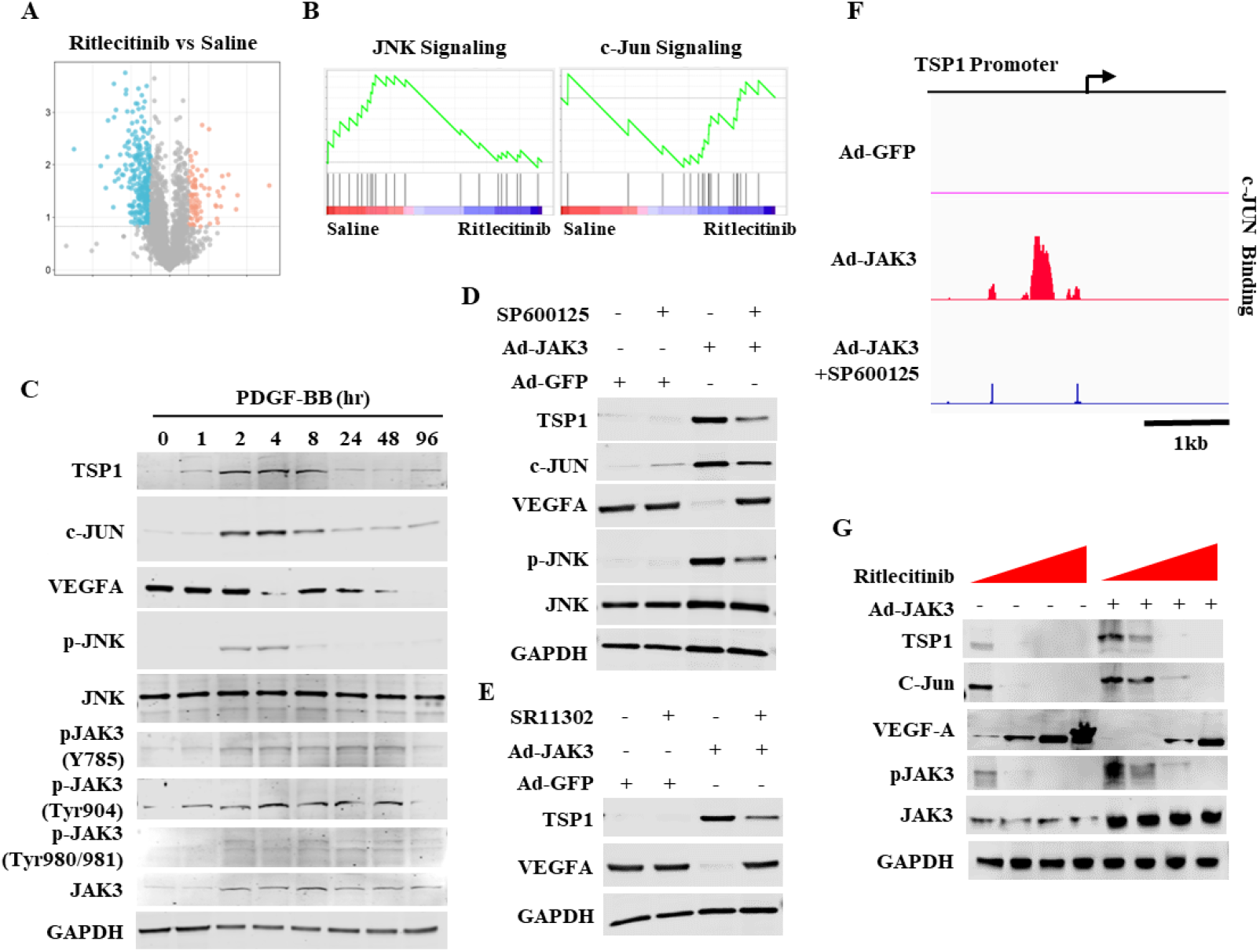
Ritlecitinib promotes VEGF-A while blocking TSP1 expression via JNK/c-Jun signaling pathways. **A-B**, ApoE-/-mouse carotid arteries were denuded by wire injury. The mice were administered with either saline or Ritlecitinib (50 mg/kg daily) starting from day 1 following injury. Carotid arteries were collected at 7 days post-injury. RNA was isolated, and RNA libraries were constructed for bulk RNA-Seq. Ritlecitinib upregulated (orange) and downregulated (blue) numerous genes as shown by Volcano plot (A). Notably, genes involved in JNK and c-Jun signaling pathways were mostly affected as shown in the enrichment score plot from GSEA (B). The green lines indicate the running enrichment score for a gene set, while the black vertical lines show the position of individual genes in the ranked gene list. The enrichment score reflects the degree to which a gene set is overrepresented at the extremes of the ranked gene list. The red and blue bars at the bottom represent gene expression correlation with the phenotype, with red indicating positive and blue indicating negative correlation. **C**, PDGF-BB time-dependently altered JAK3 and JNK phosphorylation as well as the expression of downstream genes c-Jun, VEGF-A and TSP1 in human IMA SMCs. **D**, JNK inhibitor SP600125 blocked JAK3-induced TSP1 and c-Jun expression but restored JAK3-blocked VEGF-A expression in human IMA SMCs. **E**, c-Jun inhibitor SR11302 blocked JAK3-induced TSP1 expression while restoring VEGF-A diminished by JAK3 in human IMA SMCs. **F**, JNK inhibitor SP600125 blocked JAK3-enhanced c-Jun binding to TSP1 promoter as shown by the Cut-and-Run assay. **G**, Ritlecitinib dose-dependently reversed TSP1, c-Jun, and VEGF-A expression in human IMA SMCs. Overexpression of JAK3 abolished the effects of Ritlecitinib at the low-dosages but not the high dosages, indicating minimal off-target effects of the drug.

Ritlecitinib-treated arteries, indicating that JNK and c-Jun associated transcripts were broadly suppressed by JAK3 inhibition (Fig. 5B), suggesting that JNK/c-Jun pathway, which is known to drive inflammation and SMC proliferation, might be a critical downstream mediator of JAK3 activities in vascular injury. Time-course studies showed that PDGF-BB rapidly induced JAK3 activation (phosphorylation) along with escalating JNK phosphorylation and c-Jun expression in human IMA SMCs. Notably, this was also associated with a time-dependent induction of TSP-1 and a reciprocal downregulation of VEGF-A (Fig. 5C), suggesting that active JAK3 signaling may promote TSP-1 while inhibiting VEGF-A expression in SMCs, which is unfavorable for endothelial repair. Indeed, overexpression of JAK3 increased TSP-1 and c-JUN but blocked VEGF-A expression. Pharmacologic JNK inhibition with SP600125 effectively blunted the JAK3-driven upregulation of c-Jun and TSP-1 while restored JAK3-suppressed VEGF-A expression (Fig. 5D). Similarly, specific inhibition of c-Jun activity with SR11302 abrogated JAK3-induced TSP-1 expression while rescued VEGF-A levels (Fig. 5E). These results indicated that JAK3 regulates TSP-1 and VEGF-A expression through JNK/c-Jun pathway. CUT&RUN analysis showed that JAK3 dramatically increased c-Jun binding to TSP-1 promoter, consistent with the enhanced TSP-1 expression. However, JNK inhibitor SP600125 prevented c-Jun interaction with TSP-1 promoter (Fig. 5F), further demonstrating that JAK3 promotes TSP-1 transcription by activating JNK/c-Jun cascade.

Moreover, Ritlecitinib dose-dependently modulated the expression of TSP-1, VEGF-A, and c-Jun in SMCs. Higher doses of the inhibitor led to lower c-Jun and TSP-1 levels, alongside a better restoration of VEGF-A expression (Fig. 5G). To confirm if these effects were specifically due to JAK3 blockade, we overexpressed JAK3 in Ritlecitinib-treated cells. JAK3 overexpression largely abolished the impact of low-dose Ritlecitinib on TSP-1, c-Jun, and VEGF-A by bringing their levels closer to that in untreated cells. However, higher doses of

Ritlecitinib were able to suppress c-Jun/TSP-1 while elevating VEGF-A in JAK3-overexpressing cells, suggesting that therapeutic dosages of Ritlecitinib are sufficient to block JAK3 activities with minimal off-target effect.

### JAK3 Deficiency Increases VEGF-A and Decreases TSP-1 in Injured Arteries

To determine if JAK3 regulates TSP-1 and VEGF-A expression in neointimal SMC in vivo, we examined their expression during neointima formation. As shown in Fig 6, A-B, wire injury led to a downregulation of VEGF-A and upregulation of TSP-1 protein in the arterial wall, as determined by Western blot. Remarkably, Ritlecitinib treatment reversed the injury-induced effects. Ritlecitinib restored VEGF-A protein levels and blocked the increase in TSP-1 expression (Fig 6, A-B). Co-staining of VEGF-A with ACTA2 revealed that Ritlecitinib-treated arteries had a greater proportion of SMCs expressing VEGF-A, both in the media and within the neointima (Fig. 6, C-D). Conversely, Ritlecitinib treatment dramatically reduced TSP-1 expression in SMCs of the injured arterial wall (Fig. 6E). The fraction of ACTA2+ cells that were also positive for TSP-1 was markedly lower in Ritlecitinib-treated arteries (Fig. 6F). Of note, the most pronounced changes in VEGF-A and TSP-1 expressions were observed in the neointimal regions (as indicated by the arrows in the merged images in Fig. 6, C & E), suggesting that JAK3 inhibition particularly alters neointimal SMC phenotype toward a pro-healing profile (i.e., high VEGF-A and low TSP-1 expression).

**Fig 6.**
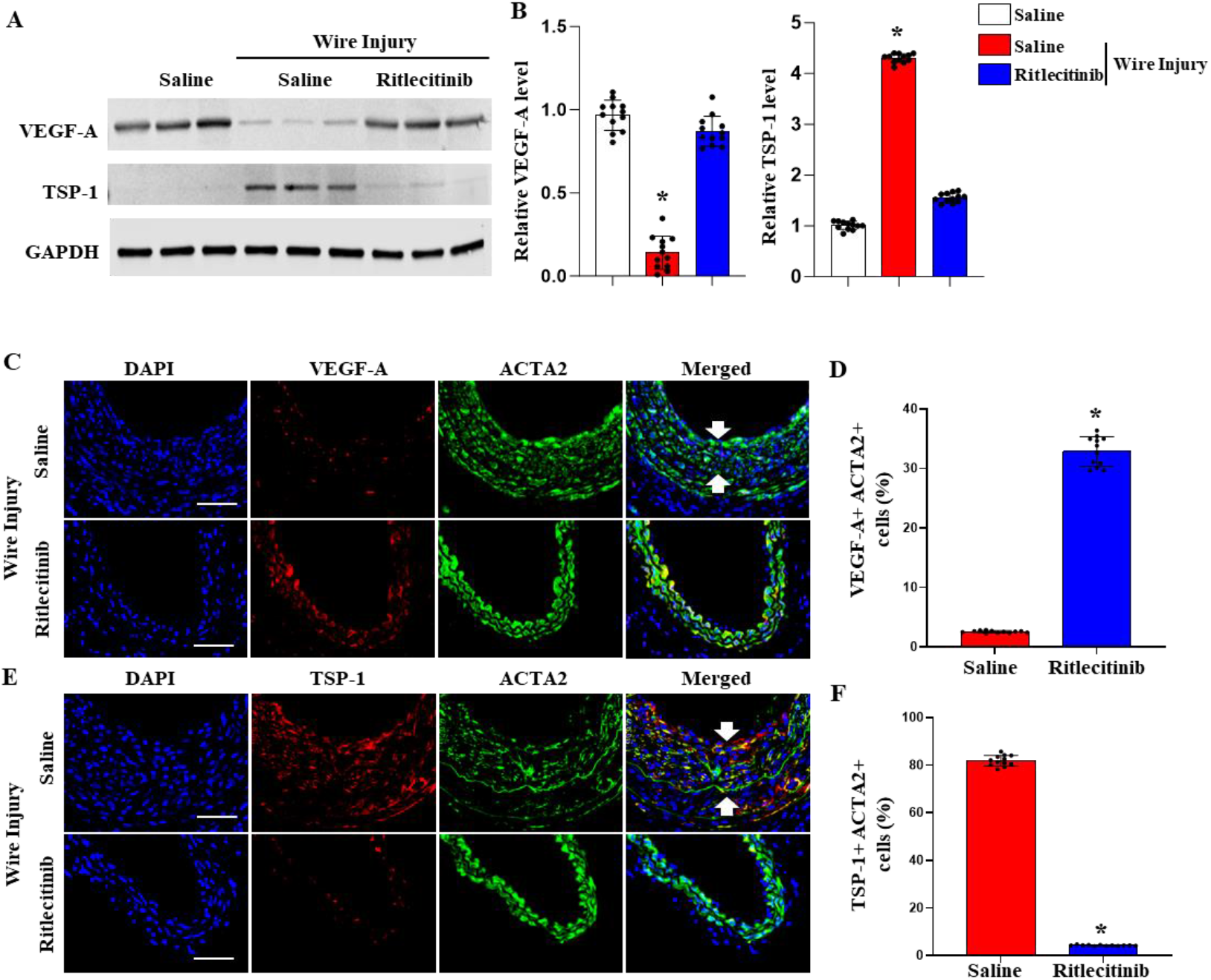
Ritlecitinib increases VEGF-A protein expression while downregulating TSP-1 in SMCs of injured arteries. **A**, ApoE-/-mouse carotid arteries were denuded by wire injury. The mice were administered with either saline or Ritlecitinib (50 mg/kg daily) starting from day 1 following the injury. The injured arteries were collected at 21 days post-injury for analysis. Ritlecitinib reversed the effects of wire injury, leading to increased VEGF-A and decreased TSP-1 expression in injured arteries, as shown by Western blot analyses. **B**, Quantification of protein expression shown in A by normalizing to GAPDH levels. P*<0.001 vs. other groups. **C-D**, Immunostaining and quantification of VEGF-A+ ACTA2+ cells (%) in saline or Ritlecitinib-treated mouse carotid arteries. Ritlecitinib significantly increased the number of VEGF-A-expressing ACTA2+ SMCs. *P<0.001 vs. saline-treated controls, n=12. **E-F**, Immunostaining and quantification of TSP-1+ ACTA2+ cells (%) in saline or Ritlecitinib-treated mouse carotid arteries. Ritlecitinib significantly decreased TSP-1 levels in ACTA2+ SMCs. *P<0.001 vs. saline-treated controls, n=12. The areas between arrows in merged images in C and E indicate neointima.

A consistent phenomenon was observed in JAK3 loss-of-function genetic models. JAK3smc-/-restored injury-suppressed VEGF-A expression while blocking the injury-induced TSP-1 expression (Supplemental Fig. 3A). Immunostaining showed that injured arteries of JAK3smc-/-mice exhibited a significantly higher percentage of VEGF-A+ SMCs in the neointima and media compared to injured WT arteries (Supplemental Fig. 3, C-D) with a substantially lower percentage of TSP-1 + SMCs (Supplemental Fig. 3, E-F). Importantly, the same results were observed in human IMAs. JAK3 knockdown via Ad-shJAK3, as well as Ritlecitinib treatment, led to higher VEGF-A and lower TSP-1 protein levels in injured human

IMA segments compared to controls, as shown by Western blot analyses (Supplemental Fig. 4). Together, these results establish that JAK3 is a crucial regulator in balancing VEGF-A and TSP-1 levels in injured arteries. By promoting TSP-1 (which can inhibit reendothelialization and promote inflammation) and suppressing VEGF-A (which supports endothelial growth and repair), JAK3 fosters conditions that favor neointimal proliferation over healing. Conversely, inhibiting JAK3 reverses this balance, i.e., upregulating VEGF-A and downregulating TSP-1, thereby enhancing endothelial regeneration and limiting neointimal expansion.

### JAK3 Regulates Neointima Formation via VEGF-A and TSP-1

To test whether the pro-healing effects of JAK3 inhibition are mediated through VEGF-A and TSP-1, we performed rescue experiments in both human arteries and mouse injury models. In the human IMA transplant model, we examined whether neutralizing VEGF-A antibody or TSP-1 protein could counteract the benefits of JAK3 knockdown. Injured IMAs were treated with Ad-shJAK3 to suppress JAK3 in the vessel wall, and then either VEGF-A neutralizing antibody (Bevacizumab) or exogenous TSP-1 protein was administered systemically one day after injury. As expected, the knockdown of JAK3 by Ad-shJAK3 significantly reduced neointima formation at 21 days post-injury. Strikingly, the therapeutic effect of JAK3 knockdown was nullified when VEGF-A was neutralized or TSP-1 was added back (Fig 7A). The neointimal area and intima/media ratio in injured IMAs with JAK3-knockdown was significantly increased by VEGF-A antibody or TSP-1 treatment (Fig. 7, B-C), demonstrating that both interventions (VEGF-A antibody or TSP-1 protein) effectively abolished the anti-neointimal benefit conferred by JAK3 silencing. These findings also indicate that enhanced VEGF-A availability and TSP-1 suppression are key mediators by which JAK3 knockdown limits neointimal hyperplasia in human arteries. A comparable outcome was observed in the JAK3smc-/-mouse model. JAK3smc-/-mice normally exhibit minimal neointima after carotid injury (Supplemental Fig 5A), as also shown earlier (Supplemental Fig. 2, A-C). However, when JAK3smc-/-mice received VEGF-A neutralizing antibody or TSP-1 protein shortly after injury, neointimal hyperplasia was largely restored. Carotid arteries of JAK3smc-/-mice treated with either agent developed thick neointimal lesions, similar to those seen in wild-type mice, effectively reversing the protective phenotype (Supplemental Fig. 5A). Morphometric analyses revealed that both the neointimal area and intima/media ratio in treated JAK3smc-/-arteries were significantly increased compared to untreated JAK3smc-/-controls (Supplemental Fig. 5, B-C).

**Fig 7.**
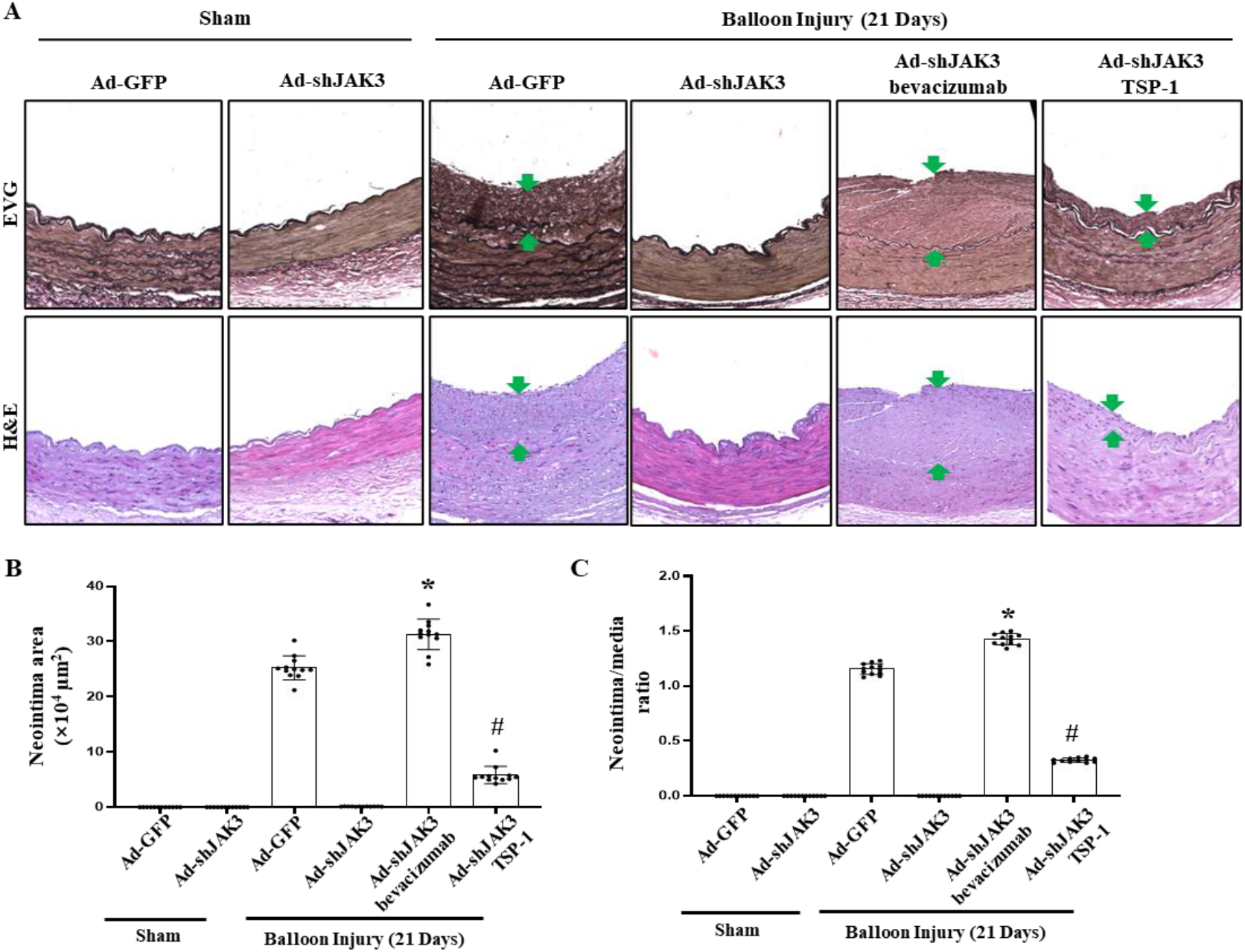
JAK3 regulates neointima formation via VEGF-A and TSP-1. Human IMA from CABG procedure were subjected to balloon injury to denude the endothelium followed by Ad-shJAK3 or Ad-GFP (control) transduction in the lumen to knock down JAK3 in artery media. The IMAs were then transplanted to abdominal aortas of immune-deficient NSG mice via end-to-end anastomosis. VEGF-A neutralizing antibody Bevacizumab (5 mg/kg) or TSP-1 recombinant protein (10 mg/kg) was administered intravenously via the tail vein on day 1 after surgery. IMAs were collected at 21 days post-injury for analyses. **A**, The effect of JAK3 knockdown on neointima hyperplasia was abolished after treatment with either VEGF-A neutralizing antibody or TSP1 protein, as shown by EVG and H&E staining. Scale bar: 100 μm. The areas between arrows indicate neointima. **B-C**, Quantification of neointima areas (B) and intima/media ratios (C) by measuring the intima and media areas. *^, #^ P<0.05 vs. the injured IMAs with Ad-shJAK3-transduction, n=12.

These results provide compelling evidence that VEGF-A and TSP-1 are critical downstream effectors of JAK3 in the vascular injury response.

### Differential Responses of Injured Human IMAs to Ritlecitinib: Implications for Precision Therapy

Although most human IMAs responded to Ritlecitinib with diminished neointimal hyperplasia following injury, we observed notable variability among individuals. Specifically, in a cohort of injured IMA specimens from different patients, most showed a significant decrease in intima/media ratio upon Ritlecitinib treatment, but a subset of arteries (5 out of 26) were non-responders, with little or no reduction in neointimal formation (Fig. 8A, red-circled data points). To uncover the basis of this differential drug response, we isolated primary SMCs from representative responders (Patients P1–P3) and non-responders (P4–P8) and analyzed their molecular responses to Ritlecitinib in vitro. We first examined the effect of Ritlecitinib on PDGF-BB-induced transcriptional activity of TSP-1 and VEGF-A in these cells using promoter luciferase reporter assays. In SMCs from the responder IMAs (P1–P3), PDGF-BB stimulation caused an expected increase in TSP1 promoter activity and a decrease in VEGF-A promoter activity. Ritlecitinib treatment effectively reversed these changes, suppressing TSP1 promoter activity while enhancing VEGF-A promoter activity (Fig. 8B), consistent with the JAK3 blockade. In contrast, SMCs from non-responders showed aberrant results. Cells from P4, P5, and P6 exhibited virtually no change in TSP1 or VEGF-A promoter activity in response to Ritlecitinib—indicating a complete lack of pathway response to the drug (Fig. 8B). Interestingly, cells from P7 and P8 displayed a partial response: Ritlecitinib was able to reduce TSP1 promoter activity in these cells, but it failed to increase VEGF-A promoter activity (Fig. 8B). This selective unresponsiveness to VEGF-A pathway in SMCs from P7/P8 hinted a distinct mechanism of resistance compared to SMCs from P4–P6.

**Fig 8.**
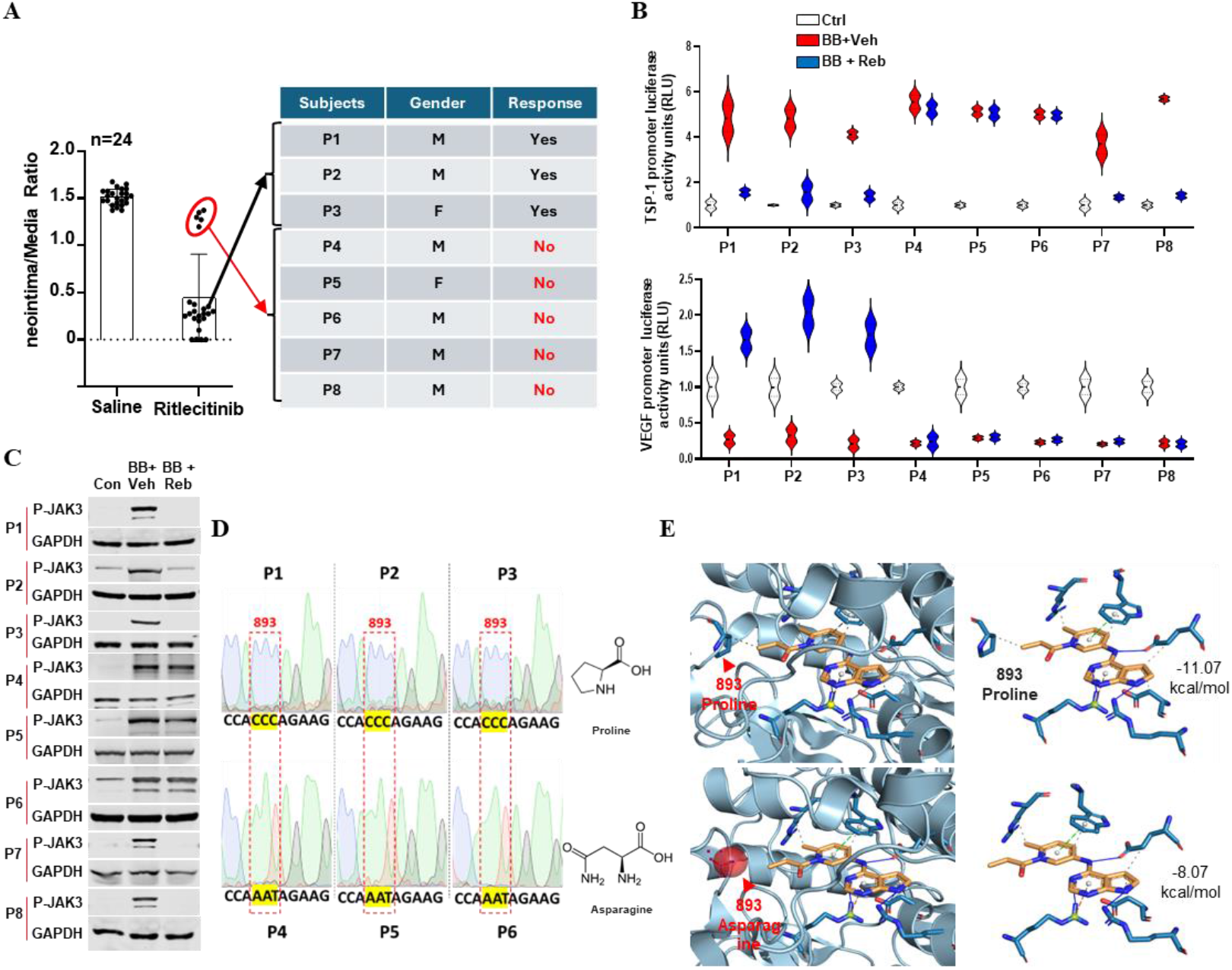
Differential responses of human IMAs with injury to Ritlecitinib treatment. **A**. differential effects of Ritlecitinib on neointima formation were observed in different IMAs with injury. Shown are intima/media ratios. Individuals circled with red represent non-responders. Three of the 19 responders (P1 to P3) and five non-responders (P4 to P8) are listed. **B**. SMCs from P1 to P8 IMAs were isolated and cultured. TSP1 and VEGF-A promoter-luciferase reporter plasmids were individually transfected into these SMCs. 24 h later, SMCs were treated with PDGF-BB with vehicle (Veh) or Ritlecitinib (Reb, 5μm) treatment for 4 h, followed by luciferase assay. Ritlecitinib reversed TSP1 and VEGF-A promoter activities altered by PDGF-BB in responders (P1-P3), but not in the non-responders (P4-P8). **C**. Ritlecitinib decreased p-JAK3 levels in SMCs of responders (P1-P3), but not in some of the non-responders (P4-P6), as analyzed by Western blotting. **D**. Sanger sequencing identified a mutation (Pro893Asn) in JAK3 protein of non-responders (red box). **E**. Molecular docking analysis revealed a significantly lower binding affinity of Ritlecitinib to the mutated JAK3 in non-responders.

By assessing JAK3 signaling or JAK3 phosphorylation in responder and non-responder SMCs, we found that Ritlecitinib treatment markedly diminished JAK3 phosphorylation levels in SMCs from responders P1–P3, confirming effective drug action on its target (Fig. 8C). However, Ritlecitinib did not reduce JAK3 phosphorylation in non-responder SMCs from P4– P6, indicating that the drug failed to inhibit JAK3 in these Cells (Fig. 8C). This finding suggested a possible alteration in the drug target itself. Indeed, Sanger sequencing of the JAK3 coding region in P4–P6 SMCs identified a point mutation resulting in a proline-to-asparagine substitution at amino acid 893 (Pro893Asn) in JAK3 kinase domain (Fig. 8D, red box). This mutation is in the ATP-binding region of JAK3 and was hypothesized to interfere with Ritlecitinib binding. Molecular docking simulations confirmed that the Pro893Asn variant significantly reduces the binding affinity of Ritlecitinib to JAK3 (Fig. 8E). Thus, a JAK3 mutation in P4–P6 likely underlies their failure to respond to Ritlecitinib, as the drug cannot effectively block the kinase domain or activity. In contrast, cells from P7 and P8 did show reduced JAK3 phosphorylation with Ritlecitinib (Fig. 8C), ruling out the target inhibition failure.

To determine the underlying cause for the lack of VEGF-A upregulation in SMCs from partial responders P7-P8, we investigated the regulatory elements of the VEGF-A gene. Sequencing analyses of VEGF-A promoter revealed that both P7 and P8 carried an identical single-nucleotide polymorphism (SNP) within a Smad2 binding motif (Supplemental Fig. 6A, red box). This SNP was absent in responders P1–P3. A CUT&RUN assay focusing on Smad2 binding to the VEGF-A promoter demonstrated that P7-P8 SMCs exhibited significantly reduced recruitment of phospho-Smad2 to the VEGF-A promoter region compared to cells with the wild-type promoter in P1-P3 SMCs (Supplemental Fig. 6B). This indicates that the SNP disrupts a transcriptional activation mechanism needed for VEGF-A induction, even when upstream signals (such as those unleashed by JAK3 inhibition) are present. Taken together, we uncovered two distinct mechanisms underlying the non-responsiveness to Ritlecitinib in human arteries: (1) a JAK3 Pro893Asn mutation that prevents Ritlecitinib from inhibiting JAK3, and (2) a VEGF-A promoter SNP that uncouples JAK3 inhibition from VEGF-A gene upregulation.

## DISCUSSION

This study demonstrates the potential of repurposing Ritlecitinib, a selective JAK3 inhibitor, as a novel systemic therapy for preventing neointimal hyperplasia while promoting endothelial recovery. Neointimal hyperplasia, a major contributor to restenosis following vascular interventions, has been historically addressed with drug-eluting stents (DES) such as those coated with paclitaxel or sirolimus^47^. While these DES have been effective in inhibiting SMC proliferation, they often fall short in promoting endothelial recovery, leading to delayed reendothelialization and subsequent risk of thrombosis and other adverse events. Our findings emphasize the importance of addressing both SMC proliferation and endothelial regeneration as part of an integrated strategy for vascular repair. Traditional anti-restenosis therapies primarily focus on inhibiting SMC proliferation to reduce neointimal growth. However, impaired reendothelialization following the suppression of SMCs can render the injured vessel segment susceptible to thrombosis and inflammation, compromising long-term vessel patency. Our results highlight that Ritlecitinib, through its dual effects of inhibiting SMC proliferation and enhancing endothelial cell proliferation, addresses the critical gap in current therapies. This dual action not only reduces neointimal hyperplasia but also accelerates the restoration of an intact endothelial layer, improving the overall outcomes of vascular interventions.

JAK3 inhibition leads to modulation of key signaling molecules involved in vascular remodeling, including VEGF-A and thrombospondin-1 (TSP-1). VEGF-A is well known for its role in promoting endothelial proliferation and migration, while TSP-1 is a potent inhibitor of angiogenesis and contributes to SMC proliferation. JAK3 inhibition by Ritlecitinib was associated with increased VEGF-A expression and decreased TSP-1 levels, thereby promoting reendothelialization while suppressing SMC-driven neointimal growth. Our data delineates a JAK3/JNK/c-Jun signaling axis that controls the expression of factors critical for vascular remodeling. JAK3 activation leads to JNK and c-Jun activation, which in turn drive up TSP-1 and downregulate VEGF-A, creating conditions that favor neointima formation and impede reendothelialization. Ritlecitinib interrupts this axis, thereby reversing these molecular effects, i.e., suppressing TSP-1 and c-Jun while boosting VEGF-A levels.

The balance between endothelial recovery and SMC inhibition is crucial for effective vascular healing and provides a distinct advantage over traditional therapies that do not adequately promote endothelial regeneration. Our study also underscores the importance of precision medicine in optimizing the use of Ritlecitinib for vascular repair. We observed variability in response to Ritlecitinib among human arteries, with some individuals exhibiting minimal therapeutic benefit. Genetic analysis identified mutations in JAK3 and SNPs in the VEGF-A promoter as underlying causes of resistance to Ritlecitinib. These findings suggest that genetic screening could be employed to identify patients who are most likely to benefit from JAK3-targeted therapy, thereby personalizing treatment and maximizing therapeutic efficacy.

Identifying such genetic variations in patients could also inform more tailored treatment strategies. For example, using alternative JAK3 inhibitors for those with resistant JAK3 mutations, or combining therapies to boost VEGF-A signaling in cases where its induction is genetically impaired.

Despite the promising findings, this study has several limitations that warrant consideration. First, while our preclinical models provide strong evidence of the efficacy of JAK3 inhibition in reducing neointimal hyperplasia and promoting endothelial recovery, the translation of these findings to clinical practice requires further validation in human trials. The safety and pharmacokinetics of Ritlecitinib in patients with cardiovascular disease must be thoroughly evaluated, especially considering its original clinical purpose for alopecia areata. Additionally, while our study provides insights into the molecular mechanisms underlying the effects of JAK3 inhibition, further research is needed to fully elucidate the downstream pathways involved and to explore potential off-target effects of Ritlecitinib in vascular system.

Future steps for clinical translation include conducting phase I/II clinical trials to assess the safety, optimal dosing, and therapeutic efficacy of Ritlecitinib in patients undergoing vascular interventions such as percutaneous coronary intervention (PCI) or coronary artery bypass grafting (CABG). These trials should also incorporate genetic screening to identify biomarkers of response, thereby facilitating a precision medicine approach to treatment. Additionally, long-term follow-up studies will be crucial to evaluate the durability of the therapeutic effects of Ritlecitinib, as well as its impact on clinical outcomes such as restenosis rates, overall cardiovascular health, and long-term vessel patency.

In conclusion, repurposing Ritlecitinib represents a promising new direction for the treatment of restenosis following vascular interventions. By simultaneously inhibiting SMC proliferation and enhancing endothelial recovery, Ritlecitinib addresses the dual challenges associated with effective vascular repair. These findings lay the groundwork for further exploration of Ritlecitinib in clinical settings, where its dual-action profile could provide substantial improvements in patient outcomes by reducing restenosis and promoting vascular healing.

## List of Supplementary Materials

Detailed Methods Supplemental Figure 1-6

## Author contributions

Conceptualization: DC, YW, SC; Methodology: DC, YW; Investigation: DC, YW; Visualization: DC, YW; Funding acquisition: DC, SC; Project administration: DC, YW, SC Clinical specimen: SL, MJ; Supervision: DC, SC; Writing – original draft: DC; Writing – review & editing: DC, SC

## Acknowledgement

We thank Ivy Sharon, Ivy Tobin, Olivia Hopkins, Lindsay Tyrer in the Surgery-Cardiothoracic team for assisting us to collect human internal mammary arteries and blood samples during the surgery operations.

## Sources of Funding

This work was supported by grants from National Institutes of Health (HL176673, HL173025, HL151444, HL119053), Department of Veterans Affairs Merit Review Awards (I01 BX006161), and American Heart Association Postdoctoral Fellowship award (#915961).

NIH public access statement: This work is the result of NIH funding, in whole or in part, and is subject to the NIH Public Access Policy. Through acceptance of this federal funding, the NIH has been given a right to make the work publicly available in PubMed Central.

## Disclosure

Authors declare that they have no competing interests

## Notes

### Competing Interest Statement

The authors have declared no competing interest.

